# Disruption of the Social Visual Pathway in Autism Spectrum Disorder

**DOI:** 10.1101/2024.09.24.614813

**Authors:** Chenhao Li, Haesoo Park, Jitendra Awasthi, Max Rolison, Mingfei Li, Dustin Scheinost, Katarzyna Chawarska, Michelle Hampson

## Abstract

The social visual pathway, which diverges from the dorsal pathway at the visual motion area (MT/V5) and runs from posterior down to anterior portions of the superior temporal sulcus (STS), specializes in processing dynamic social information. This study examined resting-state functional connectivity within this pathway in children with autism spectrum disorder (ASD) and typically developing (TD) children. Using data from the ABIDE (Autism Brain Imaging Data Exchange) repository, we found significant underconnectivity between the posterior and middle STS (pSTS-mSTS) in the right hemisphere in children with ASD compared to TD children. Weaker connectivity in this region of the pathway correlated with more severe social impairment symptoms in ASD and reduced social function across both ASD and TD groups. These findings suggest a specific disruption in the right hemisphere social visual pathway in ASD, potentially contributing to social deficits observed in the disorder.

## Background

### Disruption of the Social Visual Pathway in Autism Spectrum Disorder

Persistent deficits in social communication and interaction are the core clinical characteristics of autism spectrum disorder (ASD) [1]. Individuals with ASD often show impairments in understanding and responding to social cues, such as eye-gaze direction, facial expressions, body language, and tone of voice [2–5]. Indeed, limited attention to dynamic and speaking faces that afford eye contact represents one of the most consistently replicated biomarkers in ASD, observed during both pre-symptomatic and symptomatic stages of the disorder [3, 6–12]. While extensive research has explored these social impairments, the underlying neural mechanisms remain elusive. Recently, studies on visual processing in the brain have led to the theory of a third visual pathway specialized for social perception, which exists alongside the well-established ventral (“what”) and dorsal (“where/how”) pathways [13]. This proposed “social visual pathway” offers a novel perspective for examining the neural bases of the social deficits observed in ASD.

The social visual pathway is functionally and anatomically distinct from the ventral and dorsal pathways [13]. The traditional two-visual pathway model proposes that the ventral pathway computes object identity and the dorsal stream computes spatial and action-related information of objects [14, 15]. In contrast, the third social visual pathway specializes in processing dynamic social information, including facial and body movements, eye gaze discrimination, and the audio-visual integration of speech [13]. This pathway originates in the primary visual cortex (V1), passes through the motion-sensitive area V5/middle temporal (MT), and projects into the superior temporal sulcus (STS) [13]. Recent research has revealed a hierarchical organization within this pathway for processing social features, from low-level visual features processed in V1 and MT to high-level social communication information along the STS [16].

The STS is a key component of the social visual pathway and can be divided into three subdivisions: the posterior STS (pSTS), the middle STS (mSTS), and the anterior STS (aSTS), each with distinct roles in social perception [17, 18]. The pSTS serves as an integrative neural hub, responding to various social features and integrating social signals processed by different networks of the “social brain” [19, 20]. It has been implicated in processing dynamic faces [21, 22], biological motion [23, 24], speech sounds [25], and theory of mind [26, 27]. Moving anteriorly, the mSTS is activated by voices and auditory stimuli [18, 28, 29] as well as social reasoning [19, 30]. It shows significant responses during both auditory speech and visual speech, indicating its role in multisensory integration [31]. The aSTS has been associated with language processing, exhibiting significant responses to word- or sentence-level stimuli [18, 32]. It also shows a greater response to dynamic faces compared to static faces [21, 33] and activates during social cognition [30].

Given the critical role of the STS in social perception, it has become a region of great interest in autism research. Abnormalities in the STS have been strongly implicated in ASD, suggesting that disruptions in this region may contribute to the social deficits observed in the disorder [34–38]. Studies have consistently reported that, compared to neurotypical controls, individuals with ASD showed atypical activation patterns of the STS and its subdivisions during various social tasks, including eye gaze processing [39–41], emotion recognition [42, 43], biological motion perception [44, 45], and auditory and speech perception [46–48].

Functional connectivity studies have revealed widespread alterations in information integration between the STS, particularly the pSTS, and other brain regions involved in social cognition in ASD. For example, previous work has shown reduced functional connectivity between the pSTS and regions involved in the theory of mind network, such as the medial prefrontal cortex, during mentalizing tasks [49, 50], and between the pSTS and the fusiform face area during face processing tasks [51]. Atypical resting-state functional connectivity has also been reported between the pSTS and the inferior parietal lobule and premotor areas of the mirror neuron network [42], as well as various other social brain regions, including the amygdala, ventromedial prefrontal cortex, and fusiform gyrus [42, 52–55]. Furthermore, Alaerts et al. [56] investigated age-related changes in resting-state functional connectivity of the pSTS in individuals with ASD and found delayed and atypical developmental trajectories compared to neurotypical controls.

Despite the growing evidence of abnormalities in the STS and its functional connectivity in ASD, little is known about the functional connectivity within the social visual pathway itself, that is, between the MT/V5, pSTS, mSTS, and aSTS. Investigating the connectivity between these regions is crucial for understanding how information is processed and integrated along the social visual pathway. This gap in the literature represents an opportunity for further investigation into the neural mechanisms underlying social impairments in ASD.

In the present study, we explored whether the social visual pathway of children with ASD is disrupted as it runs down the STS. We used resting-state functional magnetic imaging (fMRI) data from the Autism Brain Imaging Data Exchange (ABIDE) repository [57, 58] to examine the resting-state functional connectivity between the nodes of this pathway in children with ASD and typically developing (TD) controls. Specifically, we investigated the resting-state functional connectivity between the MT/V5, pSTS, mSTS, and aSTS in the right and left hemispheres, respectively. Further, we explored whether alterations in resting-state functional connectivity in ASD correlate with the severity of autistic symptoms, thereby providing insights into how these disruptions may relate to behavioral manifestations of ASD. We hypothesized that children with ASD would exhibit reduced connectivity within this pathway and that lower connectivity strength would be associated with greater severity of autism symptoms and poorer social function.

## Methods and Materials

### Study Sample

The participant sample for this study was derived from our previous study [59], using data from ABIDE I and ABIDE II. The initial selection included participants aged 6-14 years from data collection sites with research-reliable clinicians administering the Autism Diagnostic Observation Schedule (ADOS) [60]. TD participants had no psychiatric diagnoses, did not use psychoactive medications, and had no family history of ASD. In addition, we included data only from sites that contributed at least 10 participants per diagnostic group with usable data. For the current analysis, we further refined the sample by excluding 12 participants due to incomplete resting-state fMRI data in areas of interest within the social visual pathway. The final dataset included 289 children diagnosed with ASD and 432 TD controls. Detailed participant characteristics are presented in Table 1.

**Table 1.** Participant Characteristics.

|  | ASD<br>(*n* = 289) | TD<br>(*n* = 432) | *P*-value |
| --- | --- | --- | --- |
| Age at scan (years, mean ± SD) | 10.4 ± 2.2 | 10.5 ± 1.6 | 0.347 |
| Sex, *n* (%) |  |  | - |
| Male | 247 (85.5) | 289 (66.9) |  |
| Female | 42 (14.5) | 143 (33.1) |  |
| Full-scale IQ (mean ± SD) | 105.2 ± 17.0 | 114.7 ± 12.5 | <0.001 |
| ADOS (module 3, raw score), *n* | 220 | - | - |
| Social Affect (mean ± SD) | 8.8 ± 3.4 | - | - |
| Restricted Repetitive Behavior (mean ± SD) | 3.1 ± 1.7 | - | - |
| SRS/SRS-2, *n* | 205 | 304 | - |
| Total Raw Score (mean ± SD) | 90.1 ± 30.0 | 19.2 ± 13.7 | <0.001 |

Both groups were matched on age, with a mean age of 10.4 years (*SD* = 2.2) in the ASD group and 10.5 years (*SD* = 1.6) in the TD group. The distribution of sex assigned at birth was 247 males and 42 females in the ASD group, compared to 289 males and 143 females in the TD group, *X*^2^ (1) = 31.30, *p* < .001. Although the groups were not matched on full-scale intelligence quotient (FIQ), all participants scored above 70. Children with ASD had a mean FIQ of 105.2 (*SD* = 17.0), whereas TD children had a mean FIQ of 114.7 (*SD* = 12.5).

To confirm the ASD diagnosis, participants with ASD were administered the Autism Diagnostic Observation Schedule-Generic (ADOS-G) [61] (178 participants) and/or the Autism Diagnostic Observation Schedule-Second Edition (ADOS-2) (110 participants) [60]. ADOS calibrated severity scores (CSS) were available for 227 participants, with a mean of 6.9 (*SD* = 1.9). Additionally, parents of 205 participants with ASD and 304 TD participants completed the Social Responsiveness Scale (SRS) [62] or Social Responsiveness Scales-2 (SRS-2) [63]. These scales measure the severity of social impairment in both individuals with ASD and the general population, with scoring compatible between editions for school-age children [64, 65]. As expected, the ASD group showed significantly higher SRS/SRS-2 raw total scores (*M* = 90.1 ± 30.0) compared to the TD group (*M* = 19.2 ± 13.7), indicating greater social impairment.

### Imaging Parameters

This study included data collected at Kennedy Krieger Institute, Georgetown University, New York University, Oregon Health & Science University, University of California Los Angeles, University of Michigan, and Yale University. Data acquisition parameters varied across sites. Details on scan parameters and site-specific protocols are available at fcon_1000.projects.nitrc.org/indi/abide/.

### Imaging Preprocessing

Motion analysis and common space registration were performed following the pipeline described in our previous study [59]. The process began with skull-stripping of anatomical images using FSL, followed by manual refinement to remove any residual non-brain tissue.

Functional data preprocessing was then carried out using SPM8. To control for potential confounding effects of excessive head motion, only participants with an average frame-to-frame displacement less than 0.15 for any run were included in the current dataset. After exclusion, in the current dataset, the ASD group showed slightly higher head motion compared to the TD group (ASD: 0.08 ± 0.03, TD: 0.07 ± 0.03, Cohen’s *d* = 0.28). The head motion was subsequently included as a covariate in analyses to minimize motion confounds.

Following motion correction, the functional images were normalized into 3mm-resolution Montreal Neurological Institute (MNI) space. This spatial normalization procedure used a previously validated algorithm [66] which combined linear registration of functional images to the skull-stripped anatomical image and subsequent nonlinear registration of the anatomical image to the MNI template brain.

### Functional Connectivity Computation

Regions of interest (ROIs) and control variables were defined using custom scripts for MATLAB. To delineate the social visual pathway, we created eight 15mm diameter spherical ROIs centered on MNI coordinates for MT/V5, pSTS, mSTS, and aSTS in both hemispheres. These coordinates were derived from Lahnakoski et al.’s [19] naturalistic study of brain activity involved in social perception. The MNI coordinates were as follows: left MT/V5 (−54, −62, 6), right MT/V5 (50, −66, −2), left pSTS (−58, −42, 12), right pSTS (58, −44, 14), left mSTS (−62, −32, 6), right mSTS (60, −22, −2), left aSTS (−56, −4, −16), and right aSTS (54, 4, −28) (see Figure 1 for a schematic representation of the regions of interest). Notably, the pSTS ROIs overlapped with the coordinates identified in Varrier and Finn’s study [67] on neural responses to social information, emphasizing their relevance to social perception.

**Figure 1.**
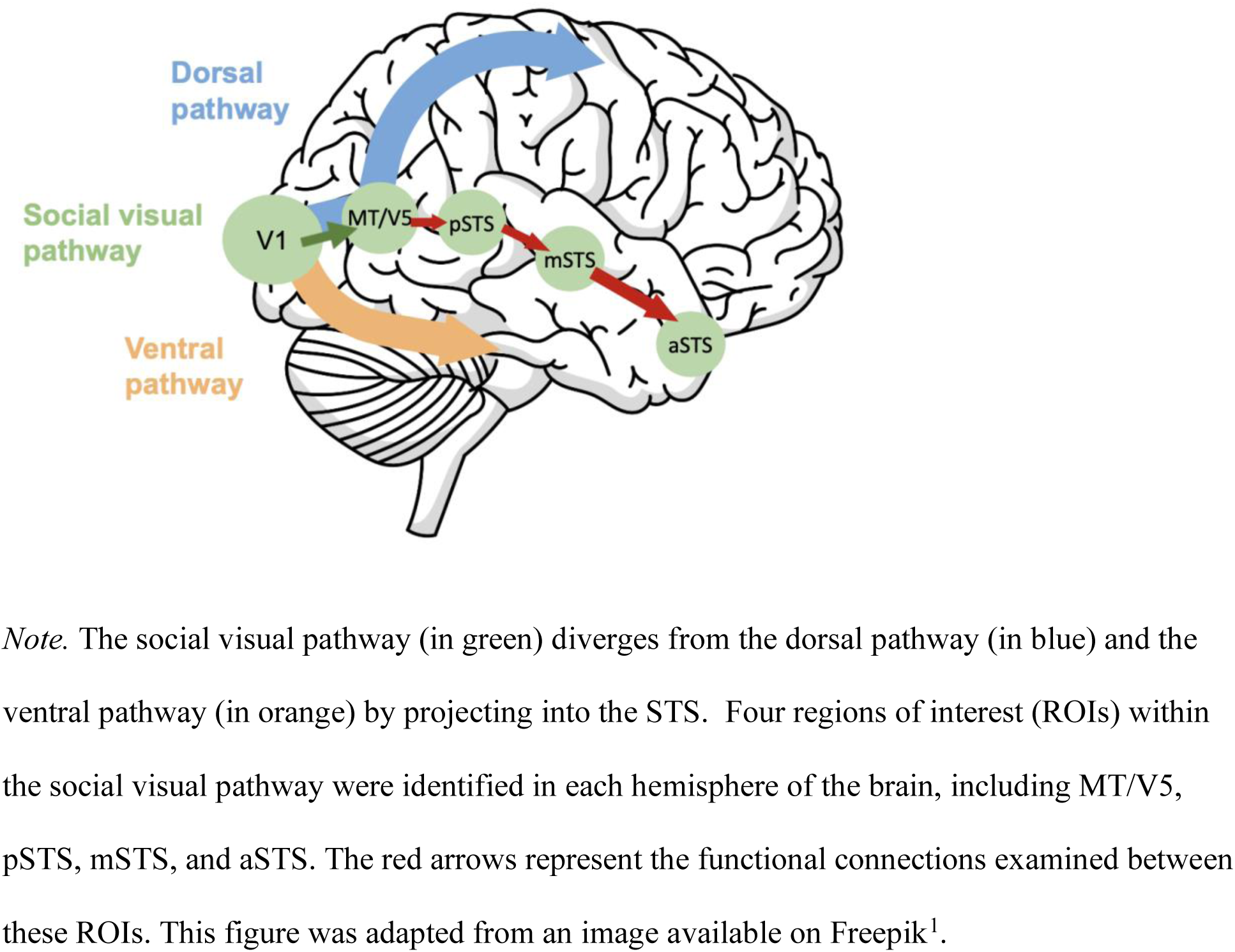
Regions of interest along the social visual pathway ^1^Image by <a href="https://www.freepik.com/free-psd/brain-outline-illustration_65105163.htm#query=brain&position=0&from_view=keyword&track=sph&uuid=56180dde-27a7-483c-b2c0-96e9d06cc841">Freepik</a>

We used partial-correlation models to compute resting-state functional connectivity between adjacent regions along the hypothesized information flow of the social visual pathway. Specifically, we examined connectivity between MT/V5 and pSTS, between pSTS and mSTS, and between mSTS and aSTS in both hemispheres. This approach was chosen to investigate how information is processed and integrated sequentially along the social visual pathway, as it runs from MT/V5 through the STS subdivisions. Time series were extracted from the voxels within each ROI from the preprocessed resting data, and the average time series across the voxels within each ROI were computed. These average time courses were then correlated using partial correlation models. To control for noise, we partialed out several nuisance factors, including linear drift, the six motion parameters from motion correction, the mean white matter signal, and the mean cerebrospinal fluid signal. The correlation coefficients were transformed to z-values using Fisher’s transformation, yielding standardized values that represent the strength of connectivity between the ROIs.

### Data Analysis

The functional connectivity strength data were analyzed using Analysis of Covariance (ANCOVA) to assess the differences in connectivity strength between ASD and TD groups while accounting for potential confounding factors. The ANCOVA models included the diagnostic group (ASD or TD) as the main predictor variable, with age, data collection site, sex, full-scale IQ, and head motion as covariates. For each participant, we computed connectivity values between three pairs of ROIs in each hemisphere. Therefore, six separate ANCOVA models were used to compare these connections across the ASD and TD groups. To address multiple comparisons across the connections examined, we applied a Bonferroni correction. The significance threshold was adjusted by dividing the alpha level of .05 by the number of tests (6), resulting in a corrected alpha level of .008. Only connections with *p*-values below this corrected threshold (.008) were considered statistically significant. All reported *p*-values were two-tailed.

In further analyses, we used generalized linear regression models to explore the association between clinical measures and the strength of any functional connections that demonstrated significant group differences. In these models, functional connectivity strength was entered as an independent variable, while measures of autism symptom severity and social impairment were used as dependent variables in separate analyses. For autism symptom severity, we used ADOS raw scores in Social Affect (SA) and Restricted and Repetitive Behaviors (RRB) domains. We exclusively used module 3 raw domain scores, calculated using the updated algorithms [68]. Participants who were administered module 2 (n = 4) or module 4 (n = 1) were excluded from these analyses. To assess social impairment across a broader spectrum, including both participants with ASD and TD participants, we examined the relationship between functional connectivity and SRS/SRS-2 raw total scores. Based on the goodness of fit of data, the Poisson model was selected for our analyses. All models included age, data collection site, sex, full-scale IQ, and head motion as covariates to reduce potential confounding effects. For SRS score analyses, we additionally included an indicator of the SRS edition to control for potential differences between the two editions of the scale.

## Results

### Connectivity Analysis

ANCOVA analyses were conducted using Bonferroni adjusted alpha levels of .008 per test (.05/6) to account for the six connections examined. Results indicated that only the functional connectivity between the pSTS and mSTS in the right hemisphere showed a significant group difference, *F* (1, 705) = 7.59, *p* = .006. Post-hoc comparison revealed that children with ASD (*M* = 0.38, *SE* = 0.02) demonstrated significantly lower connectivity between right pSTS (rpSTS) and right mSTS (rmSTS) compared to their TD peers (*M* = 0.44, *SE* = 0.02), with a mean difference of 0.06 (*t* (705) = 2.75, *SE* = 0.02) (Fig. 2). No significant group differences were found for the other five connections, at either uncorrected (*p* <.05) or Bonferroni-corrected (*p* <.008) levels. These results are also summarized in Tables 2 and 3.

**Figure 2.**
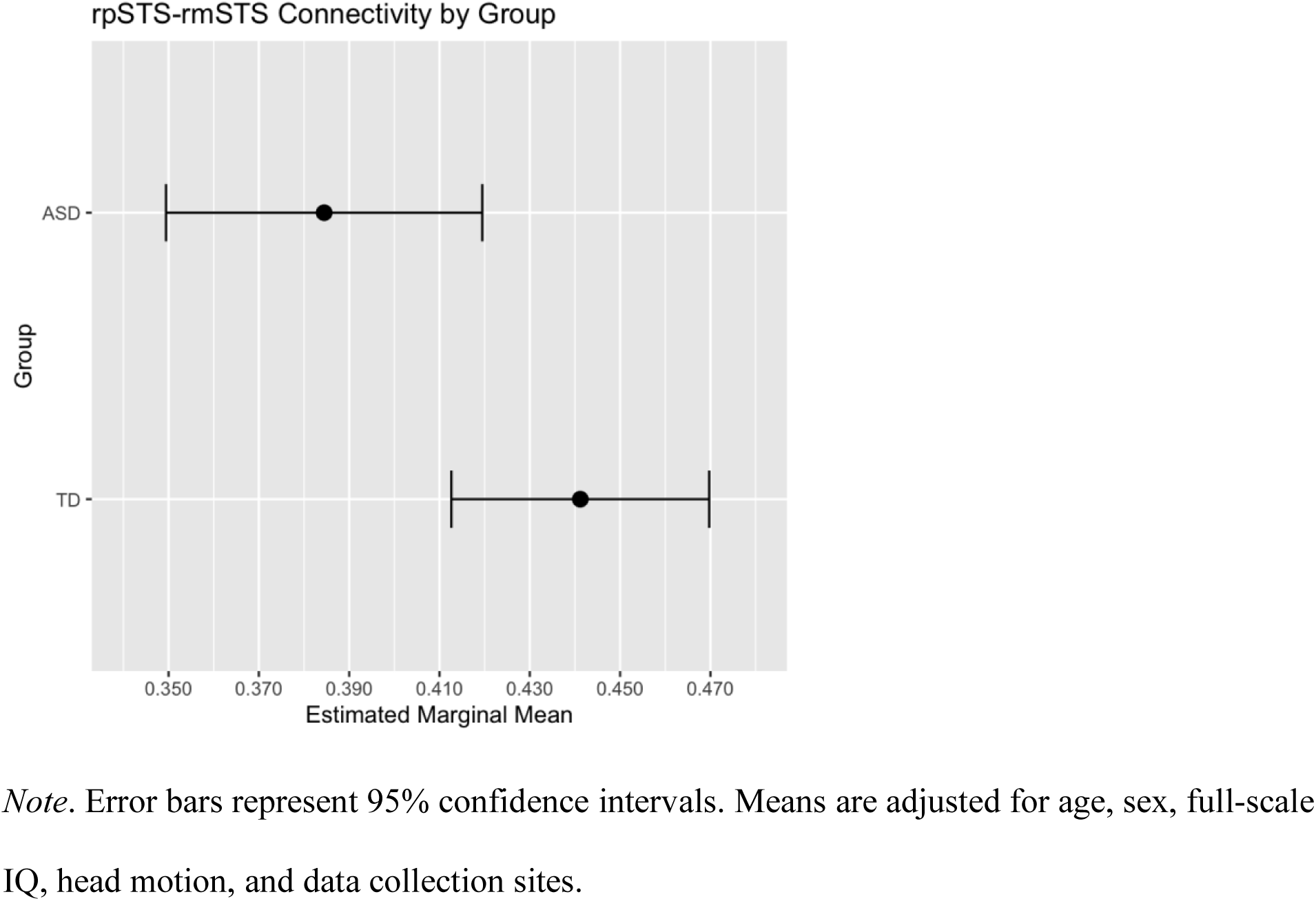
Adjusted Mean Connectivity Strength between rpSTS and rmSTS in TD and ASD Groups with 95% Confidence Intervals

**Table 2.**
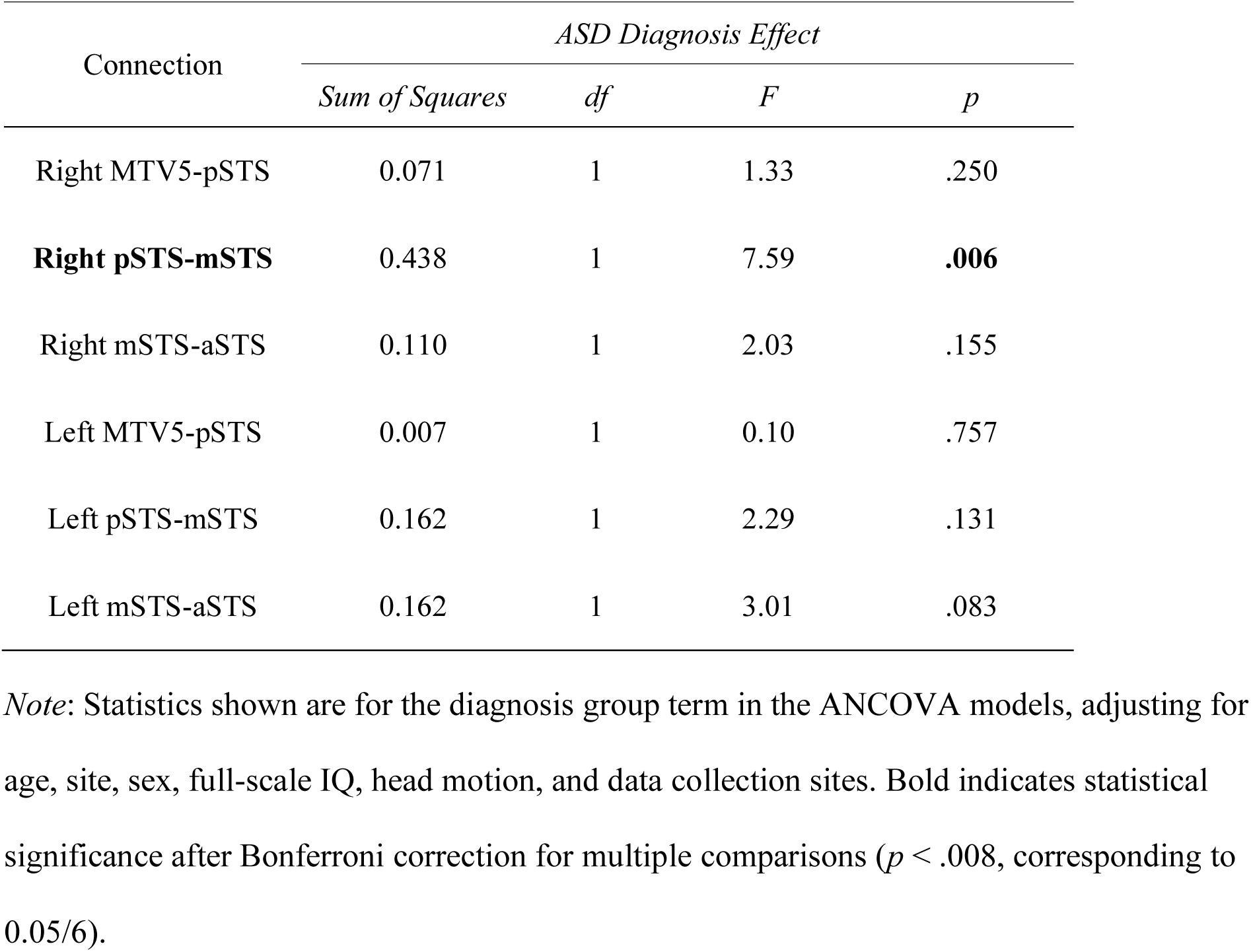
Results from six separate ANCOVA models comparing functional connectivity between ASD and TD Groups across different brain connections.

**Table 3.** Regression results for group differences in rpSTS-rmSTS connectivity.

| Predictor | $\beta$ | 95% CI | SE | *p* |
| --- | --- | --- | --- | --- |
| (Intercept) | 0.40 | [0.18, 0.61] | 0.11 | <.001 |
| **Diagnostic group (ref: TD)** | -0.06 | [-0.10, -0.02] | 0.02 | **.006** |
| Age | -0.01 | [-0.02, 0.002] | 0.01 | .103 |
| Sex (ref: female) | -0.02 | [-0.06, 0.02] | 0.02 | .371 |
| Full-scale IQ | -0.0003 | [-0.002, 0.001] | 0.001 | .607 |
| Head motion | 0.92 | [0.32, 1.53] | 0.31 | .003 |

*Note:* Site effects were included in the model but are not shown for brevity.

In an exploratory analysis, we examined the potential influence of sex on the association between ASD diagnosis and functional connectivity between the rpSTS and rmSTS. A moderation analysis revealed no significant interaction between diagnosis and sex (*p* = .801; Supplementary Table S1). Therefore, there is no evidence to suggest that the observed association between ASD diagnosis and the rpSTS-rmSTS connectivity depends on sex.

### Clinical Score Analysis

Within the ASD group, we analyzed the association with raw domain scores from ADOS. We found a significant negative association between the strength of the rpSTS-rmSTS connectivity and ADOS SA scores (*β* = −0.26, *p* = .025). However, the association between connectivity strength and ADOS RRB scores (*β* = −0.13, *p* = .50) was not significant.

To assess whether rpSTS-rmSTS connectivity strength was more broadly linked to social functioning, we analyzed SRS total raw scores in the combined ASD and TD groups. Multivariate regression analysis revealed a significant negative association between connectivity strength and SRS scores, both with (*β* = −0.09, *p* = .004) and without (*β* = −0.45, *p* < .001) the inclusion of the diagnostic group as a model indicator. These results are also summarized in Table 4.

**Table 4.**
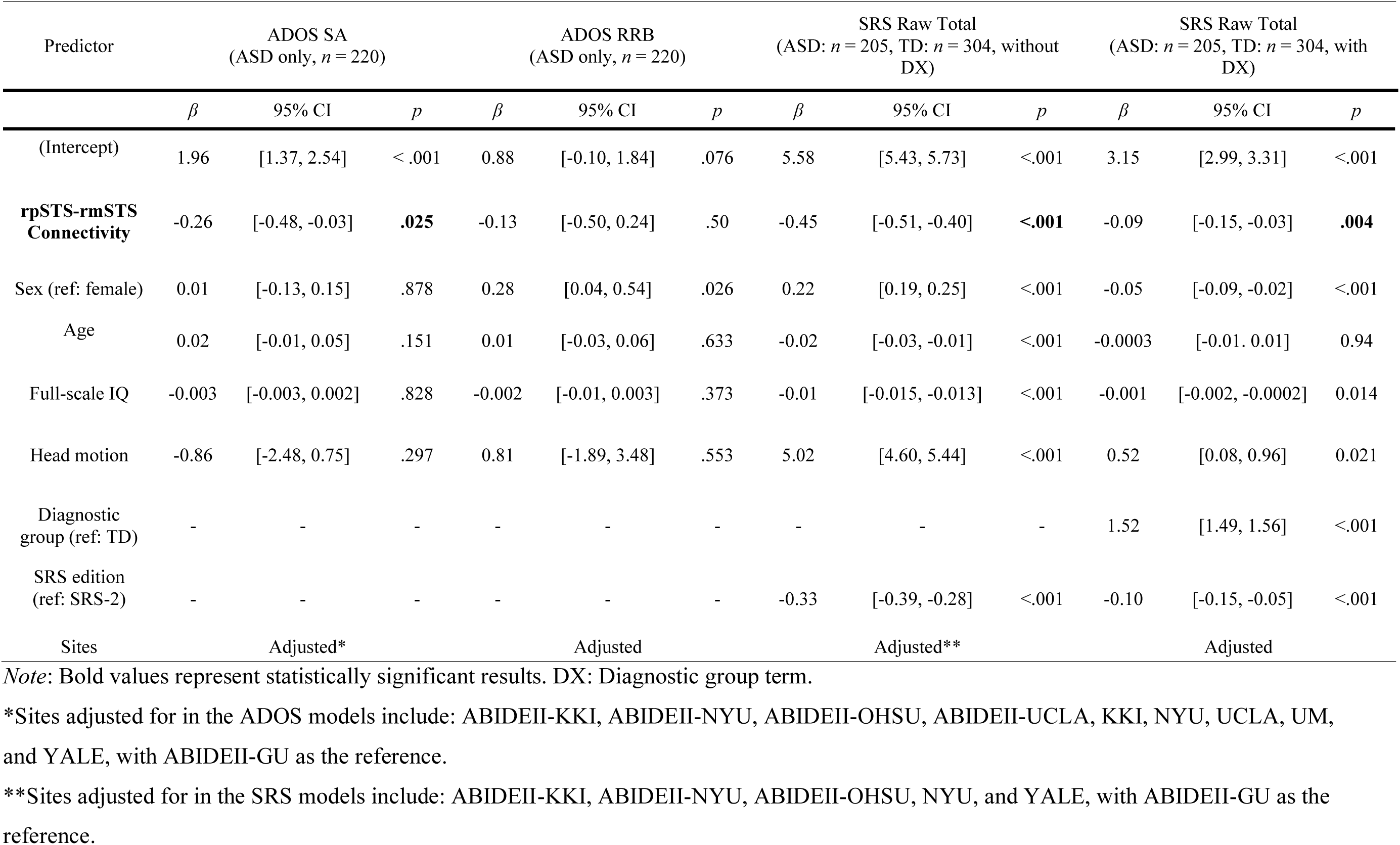
Generalized linear models evaluating associations between clinical measures and rpSTS-rmSTS connectivity strength, adjusted for sex, age, full-scale IQ, head motion, and data collection sites.

## Discussion

In this study, we used a large dataset of resting-state fMRI data from the ABIDE repository to explore potential disruptions in the social visual pathway in children with ASD. To the best of our knowledge, this is the first study to examine functional connectivity within the social visual pathway in ASD. Our findings reveal underconnectivity between the pSTS and mSTS in the right hemisphere in children with ASD compared to TD peers. We found no evidence of aberrant connectivity elsewhere along the STS, even at an uncorrected significance level, highlighting a specific region of dysfunction within the social visual pathway in ASD.

Data from a number of studies suggest that this region of the right hemisphere pathway between pSTS and mSTS is a transition zone in which the social visual pathway begins to respond to audiovisual speech stimuli [18, 19, 31]. There is also a shift in processing here from lower-level visual features to higher-level aspects of social cognition including the theory of mind [18], social interaction [19], and communication [16]. Our results are consistent with findings from eye-tracking studies examining selective social attention to visual and auditory social cues. When cues such as eye contact and speech are systematically manipulated, infants, toddlers, and school-age children with ASD exhibit marked deficits in attending to faces that require the integration of visual and speech information [3, 6–10]. Notably, these deficits are not observed when social stimuli involve dynamic but silent faces [e.g., 8]. The underconnectivity we observed between pSTS and mSTS in ASD may therefore reflect difficulties in integrating and processing complex social information, which could contribute to the social deficits observed in the disorder.

Interestingly, the disruption in functional connectivity was found only in the right hemisphere. This finding aligns with a large body of literature that has more frequently reported abnormalities in the right STS than in the left STS in individuals with ASD [e.g., 35, 40, 48, 53, 55, 56]. For example, Alaerts et al. [56] identified only pSTS in the right hemisphere as a region that could predict emotion recognition deficits in ASD and found its hypoconnectivity with the mirror network. Similarly, Pelphrey et al. [40] observed reduced connectivity between the right pSTS and the right fusiform gyrus in ASD during social cognition tasks in individuals with ASD compared to TD controls. This hemisphere-specific pattern of findings may be attributed to the functional dominance of the right STS for social communication and cognition [16, 30, 69], while the left STS is more specialized in speech and language processing [18, 29]. The specific disruption found in the right hemisphere social visual pathway in our study, consistent with previous work on right STS abnormalities in ASD, highlights the importance of the right STS in social processing and suggests that alterations in this region may contribute to social difficulties in ASD.

To further explore the role of disrupted connectivity in this part of the social visual pathway in ASD, we examined associations with clinical measures. We found that reduced connectivity between the rpSTS and rmSTS was associated with greater social impairment, both within the ASD group and more generally, across the two diagnostic groups. Specifically, within the ASD group, a significant negative relationship between the connectivity strength and ADOS SA scores indicated that weaker connectivity in this pathway is linked to greater social deficits. This supports the hypothesis that disruptions in this pathway may contribute to social impairments observed in ASD. Notably, the absence of a significant association between rpSTS-rmSTS connectivity and ADOS RRB scores underscores the specific relevance of this connection to social functioning rather than to repetitive and restricted behaviors in ASD. Furthermore, across both ASD and TD groups, we observed a significant negative association between rpSTS-rmSTS connectivity strength and SRS total raw scores, confirming that reduced connectivity is associated with more severe social challenges. Taking together, these findings highlight the importance of the rpSTS-rmSTS connection for social processing and suggest that disruptions in this portion of the pathway may be a neural mechanism underlying social deficits in ASD.

The ABIDE repository is an excellent resource for autism research, providing a large sample of neuroimaging data from children with and without ASD across a wide age range and from different research sites with varying imaging protocols. This diversity in the sample allows for the incorporation of site and age-related variability, which may reduce the signal-to-noise ratio but also yield results that are more likely to generalize across sites and ages. However, this broad sampling does not address the sex imbalance in ASD samples driven by the higher prevalence of boys with ASD [70]. We explored potential sex differences in our study but found no evidence of significant effects in our regression analysis, possibly due to the limited number of girls in the sample. While our results are suggestive, future studies with larger samples of females will be necessary to determine if our findings generalize to girls.

Furthermore, the available data in ABIDE is limited to resting-state scans. While resting-state data provides valuable insights into intrinsic functional connectivity, it does not directly capture brain activity during specific cognitive processes. Future work employing task activation studies to drive the circuitry of interest in different ways may clarify what aspects of processing within the visual pathway are most affected by rpSTS-rmSTS disruption. Moreover, experimental neurodevelopmental studies could be used to provide a finer-grained characterization of information processing deficits and their relationship to the integrity of this connection. For example, our group has previously found reduced social reward learning in children with ASD [71, 72]. The findings of our current study raise the question of whether the disrupted rpSTS-rmSTS connectivity could be related to the observed limited attention to dynamic, speaking, and expressive faces and contribute to this learning deficit. To address this question, future studies could combine social attention and social reward learning paradigms with functional neuroimaging to investigate the correlation between reward learning measures and the strength of the rpSTS-rmSTS connection. Such an approach would provide a more comprehensive understanding of the role of this portion of the social visual pathway in the social impairments observed in ASD.

The promise of neuroscience-based interventions for ASD has yet to be realized, despite their potential. Identifying neural biomarkers associated with symptoms is a crucial first step toward achieving this goal. In the present study, we report a biomarker of disconnection within the social visual pathway in ASD that is correlated with social impairment symptoms and presents a promising target biomarker for future intervention development. Correlational neuroimaging alone cannot determine whether rpSTS-rmSTS connectivity is upstream or downstream of clinical impairment. However, interventions for targeting functional brain patterns, such as fMRI neurofeedback [73], could provide an avenue for addressing these questions, and depending on the results, for developing network-based therapeutic interventions for children with ASD.

In summary, using a large dataset from the ABIDE repository, this study provides novel insights into the resting-state functional connectivity within the social visual pathway in children with ASD. Our data reveals specific underconnectivity between the pSTS and mSTS in the right hemisphere in children with ASD compared to TD peers, and the strength of this connection is associated with social function as assessed by SRS and ADOS. Together, our results suggest that this specific region of disruption within the social visual pathway in ASD is associated with social deficits. Future research should include task-based fMRI investigations and comprehensive neuropsychiatric testing to further clarify the role of the rpSTS-rmSTS connection in ASD. Identifying this specific neural connectivity pattern and understanding its relationship to social functioning is a crucial step toward developing targeted neuroscience-based interventions that could potentially improve social functioning in individuals with ASD.

## Acknowledgments

Publicly available datasets were analyzed in this study. The data can be accessed through the Autism Brain Imaging Data Exchange (ABIDE) website at http://fcon_1000.projects.nitrc.org/indi/abide/.

## References

1. American Psychiatric Association (2013): Diagnostic and statistical manual of mental disorders: DSM-5, 5th ed. Washington, D.C.: American Psychiatric Publishing.

2. Klin A, Jones W, Schultz R, Volkmar F, Cohen D (2002): Visual Fixation Patterns During Viewing of Naturalistic Social Situations as Predictors of Social Competence in Individuals With Autism. Arch Gen Psychiatry 59(9), 809–816.

3. Chawarska K, Macari S, Shic F (2012): Context modulates attention to social scenes in toddlers with autism. J Child Psychol Psychiatry 53(8), 903–913.

4. Harms MB, Martin A, Wallace GL (2010): Facial Emotion Recognition in Autism Spectrum Disorders: A Review of Behavioral and Neuroimaging Studies. Neuropsychology Review 20(3), 290–322.

5. Rutherford MD, Baron-Cohen S, Wheelwright S (2002): Reading the Mind in the Voice: A Study with Normal Adults and Adults with Asperger Syndrome and High Functioning Autism. J Autism Dev Disord 32(3), 189–194.

6. Chawarska K, Macari S, Shic F (2013): Decreased spontaneous attention to social scenes in 6-month-old infants later diagnosed with autism spectrum disorders. Biol Psychiatry 74(3), 195–203.

7. Shic F, Macari S, Chawarska K (2014): Speech Disturbs Face Scanning in 6-Month-Old Infants Who Develop Autism Spectrum Disorder. Biol Psychiatry 75(3), 231–237.

8. Shic F, Wang Q, Macari SL, Chawarska K (2020): The role of limited salience of speech in selective attention to faces in toddlers with autism spectrum disorders. J Child Psychol Psychiatry 61(4), 459–469.

9. Shic F, Naples AJ, Barney EC, Chang SA, Li B, McAllister T, et al. (2022): The Autism Biomarkers Consortium for Clinical Trials: evaluation of a battery of candidate eye-tracking biomarkers for use in autism clinical trials. Molecular Autism 13(1), 15.

10. Shic F, Barney EC, Naples AJ, Dommer KJ, Chang SA, Li B, et al. (2023): The Selective Social Attention task in children with autism spectrum disorder: Results from the Autism Biomarkers Consortium for Clinical Trials (ABC-CT) feasibility study. Autism Res 16(11), 2150–2159.

11. Macari S, Milgramm A, Reed J, Shic F, Powell KK, Macris D, Chawarska K (2021): Context-Specific Dyadic Attention Vulnerabilities During the First Year in Infants Later Developing Autism Spectrum Disorder. J Am Acad Child Adolesc Psychiatry 60(1), 166–175.

12. Vernetti A, Butler M, Banarjee C, Boxberger A, All K, Macari S, Chawarska K (2023): Face-to-face live eye-tracking in toddlers with autism: Feasibility and impact of familiarity and face covering. Autism Res.

13. Pitcher D, Ungerleider LG (2021): Evidence for a Third Visual Pathway Specialized for Social Perception. Trends Cogn Sci 25(2), 100–110.

14. Ungerleider LG, Mishkin M, Object Vision and Spatial Vision: Two Cortical Pathways, in Analysis of Visual Behavior, D.J. Ingle, M.A. Goodale, and R.J.W. Mansfield, Editors. 1982, MIT Press. p. 549–586.

15. Goodale MA, Milner AD (1992): Separate visual pathways for perception and action. Trends Neurosci 15(1), 20–25.

16. McMahon E, Bonner MF, Isik L (2023): Hierarchical organization of social action features along the lateral visual pathway. Curr Biol 33(23), 5035–5047.e5038.

17. Hein G, Knight RT (2008): Superior Temporal Sulcus—It’s My Area: Or Is It? J Cogn Neurosci 20(12), 2125–2136.

18. Deen B, Koldewyn K, Kanwisher N, Saxe R (2015): Functional Organization of Social Perception and Cognition in the Superior Temporal Sulcus. Cereb Cortex 25(11), 4596–4609.

19. Lahnakoski JM, Glerean E, Salmi J, Jääskeläinen IP, Sams M, Hari R, Nummenmaa L (2012): Naturalistic FMRI mapping reveals superior temporal sulcus as the hub for the distributed brain network for social perception. Front Hum Neurosci 6, 233.

20. Beauchamp MS, Lee KE, Argall BD, Martin A (2004): Integration of auditory and visual information about objects in superior temporal sulcus. Neuron 41(5), 809–823.

21. Pitcher D, Dilks DD, Saxe RR, Triantafyllou C, Kanwisher N (2011): Differential selectivity for dynamic versus static information in face-selective cortical regions. Neuroimage 56(4), 2356–2363.

22. Pitcher D, Duchaine B, Walsh V (2014): Combined TMS and FMRI reveal dissociable cortical pathways for dynamic and static face perception. Curr Biol 24(17), 2066–2070.

23. Allison T, Puce A, McCarthy G (2000): Social perception from visual cues: role of the STS region. Trends Cogn Sci 4(7), 267–278.

24. Morris JP, Pelphrey KA, McCarthy G (2008): Perceived causality influences brain activity evoked by biological motion. Soc Neurosci 3(1), 16–25.

25. Möttönen R, Calvert GA, Jääskeläinen IP, Matthews PM, Thesen T, Tuomainen J, Sams M (2006): Perceiving identical sounds as speech or non-speech modulates activity in the left posterior superior temporal sulcus. Neuroimage 30(2), 563–569.

26. Vander Wyk BC, Hudac CM, Carter EJ, Sobel DM, Pelphrey KA (2009): Action Understanding in the Superior Temporal Sulcus Region. Psychol Sci 20(6), 771–777.

27. Yang DY, Rosenblau G, Keifer C, Pelphrey KA (2015): An integrative neural model of social perception, action observation, and theory of mind. Neurosci Biobehav Rev 51, 263–275.

28. Belin P, Zatorre RJ, Lafaille P, Ahad P, Pike B (2000): Voice-selective areas in human auditory cortex. Nature 403(6767), 309–312.

29. Liebenthal E, Desai RH, Humphries C, Sabri M, Desai A (2014): The functional organization of the left STS: a large scale meta-analysis of PET and fMRI studies of healthy adults [Original Research]. Front Neurosci 8.

30. Dasgupta S, Tyler SC, Wicks J, Srinivasan R, Grossman ED (2017): Network Connectivity of the Right STS in Three Social Perception Localizers. J Cogn Neurosci 29(2), 221–234.

31. Venezia JH, Vaden KI, Rong F, Maddox D, Saberi K, Hickok G (2017): Auditory, Visual and Audiovisual Speech Processing Streams in Superior Temporal Sulcus [Original Research]. Front Hum Neurosci 11.

32. Humphries C, Binder JR, Medler DA, Liebenthal E (2006): Syntactic and semantic modulation of neural activity during auditory sentence comprehension. J Cogn Neurosci 18(4), 665–679.

33. Zhang H, Japee S, Stacy A, Flessert M, Ungerleider LG (2020): Anterior superior temporal sulcus is specialized for non-rigid facial motion in both monkeys and humans. Neuroimage 218, 116878.

34. Saitovitch A, Bargiacchi A, Chabane N, Brunelle F, Samson Y, Boddaert N, Zilbovicius M (2012): Social cognition and the superior temporal sulcus: Implications in autism. Revue Neurologique 168(10), 762–770.

35. Kaiser MD, Hudac CM, Shultz S, Lee SM, Cheung C, Berken AM, et al. (2010): Neural signatures of autism. Proc Natl Acad Sci 107(49), 21223–21228.

36. Sato W, Uono S (2019): The atypical social brain network in autism: advances in structural and functional MRI studies. Curr Opin Neurol 32(4).

37. Nomi JS, Uddin LQ (2015): Face processing in autism spectrum disorders: From brain regions to brain networks. Neuropsychologia 71, 201–216.

38. Zilbovicius M, Meresse I, Chabane N, Brunelle F, Samson Y, Boddaert N (2006): Autism, the superior temporal sulcus and social perception. Trends Neurosci 29(7), 359–366.

39. von dem Hagen EA, Stoyanova RS, Rowe JB, Baron-Cohen S, Calder AJ (2014): Direct gaze elicits atypical activation of the theory-of-mind network in autism spectrum conditions. Cereb Cortex 24(6), 1485–1492.

40. Pelphrey KA, Morris JP, McCarthy G (2005): Neural basis of eye gaze processing deficits in autism. Brain 128(Pt 5), 1038–1048.

41. Oberwelland E, Schilbach L, Barisic I, Krall SC, Vogeley K, Fink GR, et al. (2017): Young adolescents with autism show abnormal joint attention network: A gaze contingent fMRI study. Neuroimage Clin 14, 112–121.

42. Alaerts K, Woolley DG, Steyaert J, Di Martino A, Swinnen SP, Wenderoth N (2014): Underconnectivity of the superior temporal sulcus predicts emotion recognition deficits in autism. Soc Cogn Affect Neurosci 9(10), 1589–1600.

43. Kana RK, Patriquin MA, Black BS, Channell MM, Wicker B (2016): Altered Medial Frontal and Superior Temporal Response to Implicit Processing of Emotions in Autism. Autism Res 9(1), 55–66.

44. Freitag CM, Konrad C, Häberlen M, Kleser C, von Gontard A, Reith W, et al. (2008): Perception of biological motion in autism spectrum disorders. Neuropsychologia 46(5), 1480–1494.

45. Fourie E, Palser ER, Pokorny JJ, Neff M, Rivera SM (2019): Neural Processing and Production of Gesture in Children and Adolescents With Autism Spectrum Disorder. Front Psychol 10, 3045.

46. Boddaert N, Chabane N, Belin P, Bourgeois M, Royer V, Barthelemy C, et al. (2004): Perception of complex sounds in autism: abnormal auditory cortical processing in children. Am J Psychiatry 161(11), 2117–2120.

47. Dunham K, Zoltowski A, Feldman JI, Davis S, Rogers B, Failla MD, et al. (2023): Neural Correlates of Audiovisual Speech Processing in Autistic and Non-Autistic Youth. Multisens Res 36(3), 263–288.

48. Schelinski S, Borowiak K, von Kriegstein K (2016): Temporal voice areas exist in autism spectrum disorder but are dysfunctional for voice identity recognition. Soc Cogn Affect Neurosci 11(11), 1812–1822.

49. Moessnang C, Otto K, Bilek E, Schäfer A, Baumeister S, Hohmann S, et al. (2017): Differential responses of the dorsomedial prefrontal cortex and right posterior superior temporal sulcus to spontaneous mentalizing. Hum Brain Mapp 38(8), 3791–3803.

50. Kana RK, Keller TA, Cherkassky VL, Minshew NJ, Just MA (2009): Atypical frontal-posterior synchronization of Theory of Mind regions in autism during mental state attribution. Soc Neurosci 4(2), 135–152.

51. Weisberg J, Milleville SC, Kenworthy L, Wallace GL, Gotts SJ, Beauchamp MS, Martin A (2014): Social perception in autism spectrum disorders: impaired category selectivity for dynamic but not static images in ventral temporal cortex. Cereb Cortex 24(1), 37–48.

52. Gotts SJ, Simmons WK, Milbury LA, Wallace GL, Cox RW, Martin A (2012): Fractionation of social brain circuits in autism spectrum disorders. Brain 135(Pt 9), 2711–2725.

53. von dem Hagen EA, Stoyanova RS, Baron-Cohen S, Calder AJ (2013): Reduced functional connectivity within and between ‘social’ resting state networks in autism spectrum conditions. Soc Cogn Affect Neurosci 8(6), 694–701.

54. Abrams DA, Lynch CJ, Cheng KM, Phillips J, Supekar K, Ryali S, et al. (2013): Underconnectivity between voice-selective cortex and reward circuitry in children with autism. Proc Natl Acad Sci U S A 110(29), 12060–12065.

55. Lee JK, Amaral DG, Solomon M, Rogers SJ, Ozonoff S, Nordahl CW (2020): Sex Differences in the Amygdala Resting-State Connectome of Children With Autism Spectrum Disorder. Biol Psychiatry Cogn Neurosci Neuroimaging 5(3), 320–329.

56. Alaerts K, Nayar K, Kelly C, Raithel J, Milham MP, Di Martino A (2015): Age-related changes in intrinsic function of the superior temporal sulcus in autism spectrum disorders. Soc Cogn Affect Neurosci 10(10), 1413–1423.

57. Di Martino A, Yan CG, Li Q, Denio E, Castellanos FX, Alaerts K, et al. (2014): The autism brain imaging data exchange: towards a large-scale evaluation of the intrinsic brain architecture in autism. Mol Psychiatry 19(6), 659–667.

58. Di Martino A, O’Connor D, Chen B, Alaerts K, Anderson JS, Assaf M, et al. (2017): Enhancing studies of the connectome in autism using the autism brain imaging data exchange II. Scientific Data 4(1), 170010.

59. Rolison M, Lacadie C, Chawarska K, Spann M, Scheinost D (2021): Atypical Intrinsic Hemispheric Interaction Associated with Autism Spectrum Disorder Is Present within the First Year of Life. Cereb Cortex 32(6), 1212–1222.

60. Lord C, Rutter M, DiLavore PC, Risi S, Gotham K, Bishop S, Autism Diagnostic Observation Schedule, Second Edition. 2012, Western Psychological Services: Torrance, CA.

61. Lord C, Rutter M, DiLavore PC, Risi S, Autism diagnostic observation schedule—WPS (ADOS-WPS). 2002, Los Angeles (CA): Western Psychological Services.

62. Constantino JN, Gruber CP, Social Responsiveness Scale (SRS). 2007, Western Psychological Services: Los Angeles, CA.

63. Constantino JN, Gruber CP, Social Responsiveness Scale, Second Edition (SRS-2). 2012, Western Psychological Services: Torrance, CA.

64. Frazier TW, Ratliff KR, Gruber C, Zhang Y, Law PA, Constantino JN (2013): Confirmatory factor analytic structure and measurement invariance of quantitative autistic traits measured by the Social Responsiveness Scale-2. Autism 18(1), 31–44.

65. Bruni TP (2014): Test Review: Social Responsiveness Scale–Second Edition (SRS-2). Journal of Psychoeducational Assessment 32(4), 365–369.

66. Scheinost D, Kwon SH, Lacadie C, Vohr BR, Schneider KC, Papademetris X, et al. (2017): Alterations in Anatomical Covariance in the Prematurely Born. Cereb Cortex 27(1), 534–543.

67. Varrier RS, Finn ES (2022): Seeing Social: A Neural Signature for Conscious Perception of Social Interactions. J Neurosci 42(49), 9211–9226.

68. Gotham K, Risi S, Pickles A, Lord C (2007): The Autism Diagnostic Observation Schedule: revised algorithms for improved diagnostic validity. J Autism Dev Disord 37(4), 613–627.

69. Sliwinska MW, Pitcher D (2018): TMS demonstrates that both right and left superior temporal sulci are important for facial expression recognition. NeuroImage 183, 394–400.

70. Maenner MJ, Warren Z, Williams AR, Amoakohene E, Bakian AV, Bilder DA, et al. (2023): Prevalence and Characteristics of Autism Spectrum Disorder Among Children Aged 8 Years - Autism and Developmental Disabilities Monitoring Network, 11 Sites, United States, 2020. MMWR Surveill Summ 72(2), 1–14.

71. Wang Q, Chang J, Chawarska K (2020): Atypical Value-Driven Selective Attention in Young Children With Autism Spectrum Disorder. JAMA Netw Open 3(5), e204928.

72. Wang Q, DiNicola L, Heymann P, Hampson M, Chawarska K (2018): Impaired Value Learning for Faces in Preschoolers With Autism Spectrum Disorder. J Am Acad Child Adolesc Psychiatry 57(1), 33–40.

73. Hampson M, ed. fMRI neurofeedback. (2021), London: Academic Press, Elsevier.

