## Supplementary Table S1 for "Disruption of the Social Visual Pathway in Autism Spectrum Disorder"

*Regression analysis of sex effects on rpSTS-rmSTS functional connectivity*

| Predictor | *β* | 95% CI | *SE* | *p* |
| --- | --- | --- | --- | --- |
| (Intercept) | 0.40 | [0.18, 0.61] | 0.11 | <.001 |
| Diagnostic group  (ref: TD) | -0.07 | [-0.15, -0.02] | 0.02 | .129 |
| Age | -0.01 | [-0.02, 0.002] | 0.01 | .102 |
| Sex (ref: female) | -0.02 | [-0.07, 0.03] | 0.02 | .370 |
| Full-scale IQ | -0.0003 | [-0.002, 0.001] | 0.001 | .601 |
| Head motion | 0.93 | [0.32, 1.53] | 0.31 | .003 |
| Diagnostic group * Sex | 0.01 | [-0.08, 0.11] | 0.05 | 0.80 |

*Note*: Site effects were included in the model but are not shown for brevity.
